# A kinetic error filtering mechanism for enzyme-free copying of nucleic acid sequences

**DOI:** 10.1101/2021.08.06.455386

**Authors:** Tobias Göppel, Benedikt Obermayer, Irene A. Chen, Ulrich Gerland

## Abstract

Accurate copying of nucleic acid sequences is essential for self-replicating systems. Modern cells achieve error ratios as low as 10^-9^ with sophisticated enzymes capable of kinetic proofreading. In contrast, experiments probing enzyme-free copying of RNA and DNA as potential prebiotic replication processes find error ratios on the order of 10%. Given this low intrinsic copying fidelity, plausible scenarios for the spontaneous emergence of molecular evolution require an accuracy-enhancing mechanism. Here, we study a ‘kinetic error filtering’ scenario that dramatically boosts the likelihood of producing exact copies of nucleic acid sequences. The mechanism exploits the observation that initial errors in template-directed polymerization of both DNA and RNA are likely to trigger a cascade of consecutive errors and significantly stall downstream extension. We incorporate these characteristics into a mathematical model with experimentally estimated parameters, and leverage this model to probe to what extent accurate and faulty polymerization products can be kinetically discriminated. While limiting the time window for polymerization prevents completion of erroneous strands, resulting in a pool in which full-length products show an enhanced accuracy, this comes at the price of a concomitant reduction in yield. We show that this fidelity-yield trade-off can be circumvented via repeated copying attempts in cyclically varying environments such as the temperature cycles occurring naturally in the vicinity of hydrothermal systems. This setting could produce exact copies of sequences as long as 50mers within their lifetime, facilitating the emergence and maintenance of catalytically active oligonucleotides.

## INTRODUCTION

Accurate copying of genetic information is essential for the emergence of life [1–3]. In extant cells, template-directed polymerization of polynucleotides is catalyzed by sophisticated enzymatic machineries, which mitigate and correct copying errors [4]. A key enzymatic mechanism is kinetic proofreading, an on-the-fly correction scheme that uses chemical energy to perform multiple discrimination steps between correct and incorrect nucleotides [5–7], enabling remarkably low error ratios, e.g., between 10^-10^ and 10^-8^ per base pair for DNA replication. However, life must have emerged without complex enzymes [8–10].

Enzyme-free copying of short information-carrying polymers such as RNA or DNA strands has been studied extensively [11, 12]. In particular, non-enzymatic template-directed polymerization has become an established experimental model system to investigate prebiotic modes of copying: A short strand bound to a longer ‘template’ strand is sequentially extended at its 3’-end with single nucleotides or short oligomers [13–18], producing a (partial) complementary copy of the template. Lacking an inherent correction mechanism, errors during this copying process are frequent [14, 19–21]. Experiments in the presence of all four bases suggest that not even genetic information as short as ten nucleotides could be maintained by non-enzymatic template-directed polymerization [22].

How could accurate copying of genetic information be achieved without complex enzymes? A possible precursor to kinetic proofreading, which actively corrects errors right after they occur (Fig. 1A), is a passive error filtering mechanism, in which erroneous copies are not corrected, but preferentially eliminated or separated based on their physicochemical properties. How could such error filtering arise in a prebiotically plausible scenario? A key experimental observation is that the speed of template-directed polymerization strongly depends on the sequence context [15, 19, 23]. Mismatches at the 3’-terminus of a partial copy slow down the extension reaction by one or two orders of magnitude [20], and facilitate the incorporation of further non-complementary nucleotides, leading to error clusters [21]. The first effect, called post-mismatch stalling, was originally discovered in enzymatic copying before it was observed in enzyme-free systems [24]. In combination, post-mismatch stalling and error clustering cause erroneous partial copies to grow slowly, opening the door to an error filtering mechanism based on kinetic discrimination (Fig. 1B): With a limited time window for copying, only copies with no or few errors can reach full length, such that any physicochemical process that is length-selective can achieve error filtering (Fig. 2A).

**FIG. 1.**
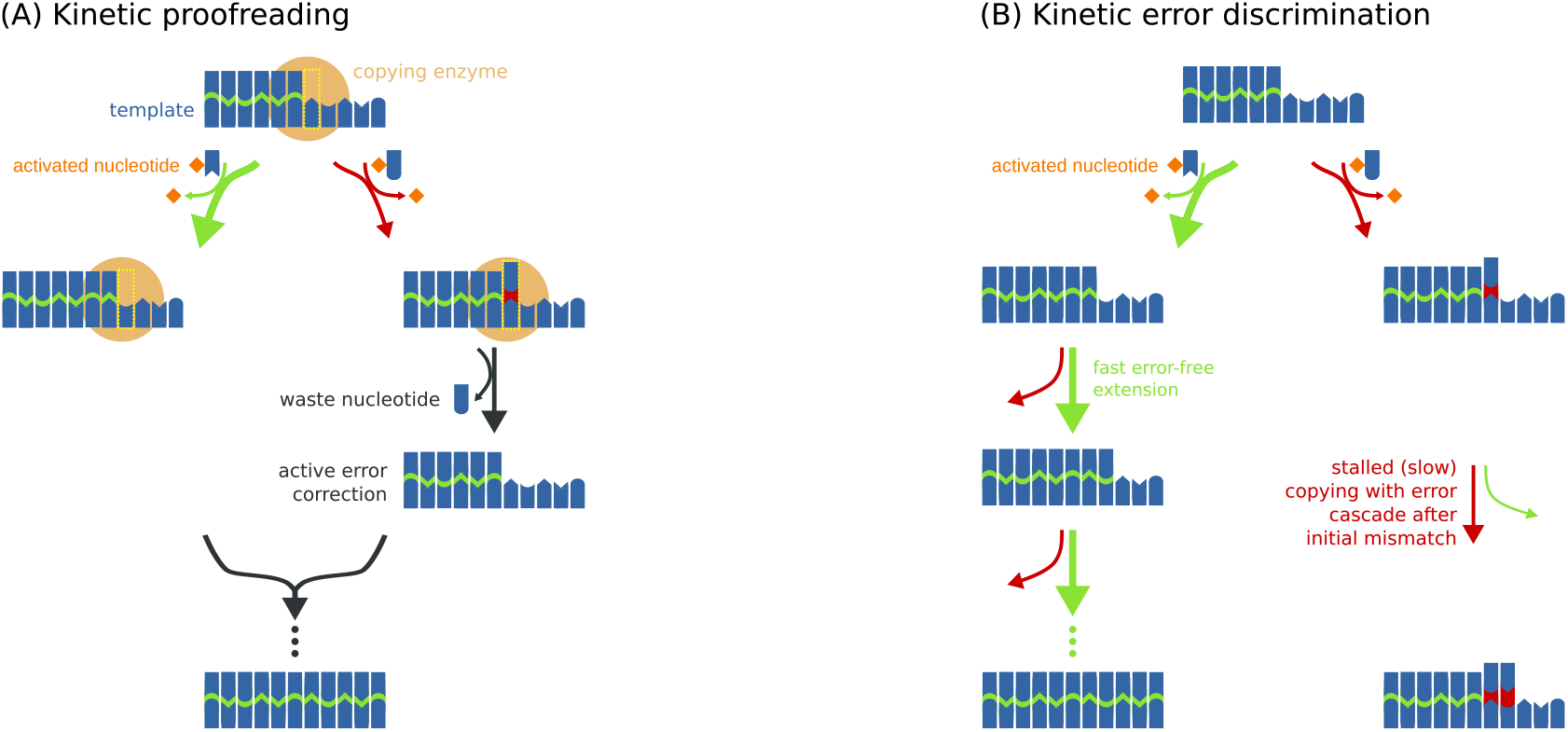
Kinetic proofreading versus kinetic error discrimination. (A) Kinetic proofreading adds an error correction step on top of the thermodynamic discrimination between correct and incorrect nucleotides. A polymerase with proofreading ability can remove a covalenty attached mismatch, allowing for a second chance to incorporate the correct nucleotide. The error correction is coupled to the consumption of chemical energy. (B) In contrast, errors occurring during non-enzymatic copying remain. The accuracy is controlled by only one discrimination step, and cannot exceed the thermodynamic limit set by the intrinsic discrimination free energy. However, an initial error typically triggers a cascade of consecutive errors and kinetically stalls the speed of downstream extension, allowing for kinetic error discrimination: If the polymerization process is stopped after a limited time, accurate copies reach full length, whereas erroneous strands remain as short waste products.

**FIG. 2.**
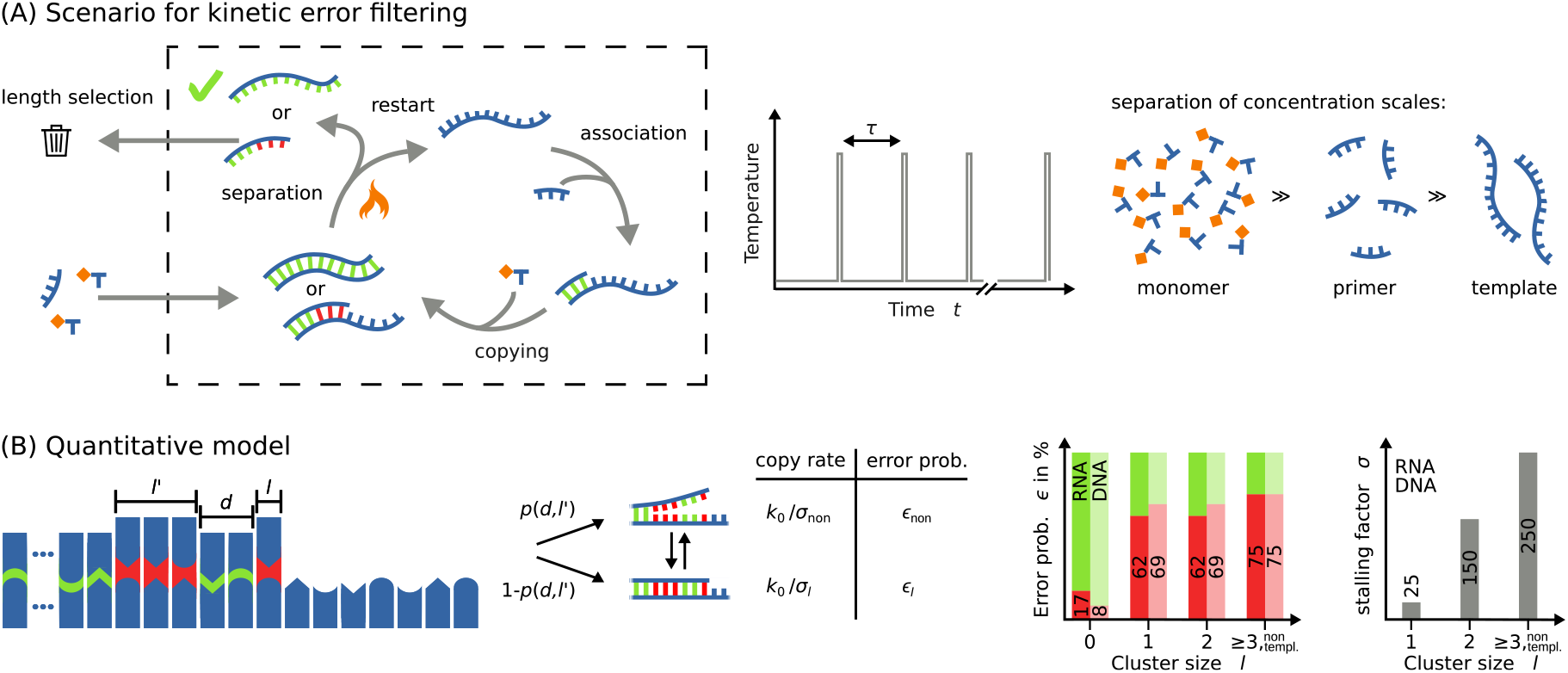
Scenario and model for kinetic error filtering. (A) To support kinetic error filtering, a suitable environment must provide limited time windows for non-enzymatic copying and length-selective transport or adhesion of polynucleotides. We envisage a scenario, in which e.g. the ambient temperature changes periodically, leaving only time windows of typical duration *τ* for the association between complementary strands and template-directed polymerization. Furthermore, we posit that the system leaks shorter oligonucleotides, and thereby preferentially removes erroneous copies. Conversely, monomers and short ‘primer’ oligonucleotides can also readily enter the system from the external environment. (B) Our quantitative model for non-enzymatic copying distinguishes only between matching and mismatching base pairs. The length *l* of an error cluster at the extension site determines the stalling factor *σ_l_* and the error fraction *ϵ_l_* (numerical values shown in the bar plots). Additionally, the distance *d* to the preceding error cluster and its size *l*′ affect the extension mode: The next extension occurs at the reduced speed and fidelity of a non-templated process with the probability *p*(*d*, *l*′) for an unbound terminus.

The two prerequisites for kinetic error filtering can both be provided by non-equilibrium environments, e.g., on the early Earth: (i) A limited time window for the copying process emerges when the ambient temperature, pH, or molecular concentrations change periodically, from conditions that promote base-pairing to conditions that favor dissociation of hybridized strands [25–30]. (ii) Length-selective physical properties, e.g. transport in thermal gradients [31, 32], accumulation on mineral surfaces [33], or retention within lipid vesicles [34], can cause a preferential loss of shorter strands. Both of these conditions can be simultaneously met in hydrothermal systems [29, 30, 32].

If kinetic error filtering is a plausible accuracy-enhancing mechanism, by how much could it boost the accuracy? Would it not reduce the yield of the copying process such as to annihilate its beneficial effects? And, most importantly, could kinetic error filtering be sufficiently effective to support the spontaneous emergence and maintenance of catalytically active oligonucleotides by template-directed polymerization? These questions intrinsically require a quantitative analysis, which we provide here.

Prior work [20] studied the beneficial effect of post-mismatch stalling on Eigen’s error threshold [1], within a coarse-grained mutation-selection model of two replicators competing in an environment with constant carrying capacity. In contrast, we consider a primordial scenario, in which the accuracy and yield of a primitive copying process must be sufficient to form at least one accurate copy for a template, on average, before the template is destroyed, e.g., by hydrolysis [15]. If this condition is not met, then any accidental discovery of a weakly catalytic sequence by random assembly will be lost again before it can further evolve. We base our analysis on a quantitative model rooted in data from primer-extension experiments with DNA and RNA, including also the effect of error clustering [21]. Using this model, we explicitly study the stochastic kinetics of template-directed polymerization in cyclic environments that offer only limited time windows for polymerization. We first characterize the fidelity-yield trade-off that emerges within a single such time window. Our subsequent analysis then reveals that cyclic environments can effectively break this fidelity-yield trade-off. This permits kinetic error filtering to facilitate the emergence and maintenance of catalytically active oligonucleotides, by significantly increasing the sequence length for which correct copies can be obtained within the lifetime of a template.

## RESULTS

### Kinetic error filtering versus kinetic proofreading

Correcting errors right after they occur is a natural solution to the problem of high intrinsic error rates. The evolutionary origin of kinetic proofreading is not clear, but extant cells use this principle not only in their processive copying enzymes mediating transcription [4], replication [35], and translation [36], but also in non-processive enzymes such as tRNA synthetase [7] or the T-cell receptor complex [37]. However, even a minimal, non-processive copying scheme with proofreading would require an enzyme that can perform a conformational transition coupled to energy release, in addition to catalyzing backbone bond formation (Fig. 1A). Therefore, it appears highly unlikely that a gratuitous proofreading mechanism would be available to primitive prebiotic replicating systems.

Without error correction, errors escaping the intrinsic thermodynamic discrimination will remain, unless erroneous copies are preferentially removed from the system. Kinetic error filtering is a two-step mechanism for such a preferential removal: First, it kinetically suppresses the formation of erroneous full length copies by limiting the copying process to a finite time window in a cyclic environment. Second, it preferentially leaks shorter strands out of the system, thereby removing the strands that contain most errors. From a thermodynamic perspective, kinetic error filtering is driven by (a part of) the free energy dissipated in the environment. This is in contrast to kinetic proofreading, where an enzyme couples the dissipation of chemical energy to error correction [5, 6].

To function, kinetic error filtering requires a suitable non-equilibrium environment. We will consider a scenario of the type illustrated in Fig. 2A: A leaky compartment is embedded in an aqueous environment providing a mixture of chemically reactive nucleotides and their polymerization products. Longer sequences have an increased residence time within this compartment, due to their charge or physical size. We do not make any assumption about the specific mechanism mediating the retention of longer sequences; it could be based on surface interactions [38], size-dependent transport through the compartment boundary [34], or bulk transport effects such as thermophoresis and convection [32, 39]. While spontaneous polymerization produces oligonucleotides with a statistical distribution of chain lengths [40], with higher-order oligomers much less (in general, exponentially less [41]) likely than monomers, stochastic fluctuations may occasionally lead to a long ‘template’ sequence within such a compartment. We assume that the physicochemical conditions (temperature, pH, or salt concentrations) in the vicinity of a template display a cyclic variation, such that the hybridization of short oligomers acting as ‘primer’ sequences only occurs within time windows of typical duration τ. The cyclic variation may arise from internal convection cycles [30] or from external periodic variations. To be more concrete, we will consider the case of temperature cycles for our model. The key assumption is that (partial) copies separate from their template at the end of a cycle, and that the probability for rebinding in the next cycle is low, since short primer molecules are much more abundant.

A submerged porous rock exposed to a temperature gradient could provide one natural realization of a suitable non-equilibrium environment. Within a pore, the interplay of convection and thermophoresis leads to a flow field, in which molecules move and experience periodic temperature changes, as has been demonstrated experimentally with a controlled lab setup [30]. In this case, polynucleotides of length 35 experienced temperature cycles featuring a short peak, during which double strands dehybridize. The copying time windows *τ* within such thermal flow chambers are controlled by the chamber geometry [31].

In the quantitative analysis presented next, we will see that efficient kinetic error filtering imposes conditions on the timescale *τ* depending on the template length *L*. One might then object that this scenario for kinetic error filtering requires “fine-tuning” of conditions. However, natural compartments and pores come in a broad range of sizes and geometries, and this natural variation can produce a correspondingly broad range of timescales *τ*. As a consequence, the massively parallel nature of natural experiments eliminates the potential fine-tuning issue.

### Template-directed polymerization in a limited time

To analyze kinetic error discrimination within a finite time *τ*, we use a mathematical model based on experimental characterizations of non-enzymatic template-directed polymerization [14, 20, 21]. Within this model, template-directed integration of monomers proceeds at a basal extension rate *k*_0_ in the absence of any mismatches. This rate defines the basal extension timescale *t*_0_ = 1/*k*_0_, which serves as the elementary time unit for this study, since the actual experimental timescale depends on the precise chemical conditions, including the type of leaving group used for the chemical activation of nucleotides [42]. In typical experiments, *t*_0_ is on the order of one hour.

We parameterize the probability *ϵ* for a copying error and the stalling factor *σ* as a function of the local structure of the template-copy complex at the extension site (Fig. 2B). This structure is described by (i) the number *l* ≥ 0 of successive mismatches directly at the extension site, (ii) the size *l*′ ≥ 0 of the next error cluster further upstream of the extension site, and (iii) the distance *d* > 0 to this next error cluster. Based on the values *d* and *l*′, we estimate the probability *p*(*d*, *l*′) that a terminus following a series of mismatches is in an unbound dangling-off configuration [21], see *Materials and methods*.

A dangling terminus is extended with the error probability of an unbiased, non-templated extension, *ϵ*_non_ = 0.75, and the corresponding extension rate is reduced by the stalling factor *σ*_non_ = 250 to *k*_0_/*σ*_non_ [21, 43]. If the terminus is closed, i.e., with probability 1 – *p*(*d*, *l*′), the stalling factor *σ_l_* and the error probability *ϵ_l_* depend only on the number *l* of mismatches at the extension site. The associated copying rate is

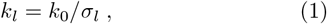

where *σ*_0_ = 1 and the values for *l* > 0 are given in Fig. 2B. Experimentally, the basal extension rate, the stalling factors, and the error probabilities also depend on the exact sequence context, i.e, the templating and the incoming nucleotide as well as their neighbors [15, 17, 19, 20, 22, 23, 44]. However, averaging over results obtained for many random sequences with sequence-dependent parameters was found to be essentially equivalent to using sequence-averaged parameters instead [21]. This justifies the practical simplification that our model makes by distinguishing only between matching and nonmatching base pairs. The average error probability *ϵ*_0_ for extending a closed terminus with no mismatch is 0.08 for DNA and 0.17 for RNA [14, 45]. After a first mismatch, the error probability increases more than sevenfold for DNA and roughly threefold for RNA [21]. Systematic measurements of the copying accuracy following error clusters of size two are missing, but the existing data suggest that the error probability remains unchanged [21]. Typically, an initial mismatch stalls the copying speed by one to two orders of magnitude. A second mismatch then slows down the extension by another factor of six [20, 21].

This mathematical model corresponds to a Markov process, in which the stochastic template-directed polymerization dynamics depend only on the current state of the terminus. We analyze these dynamics with simulations based on the Gillespie algorithm [46] and with analytical approximations described below. For an overview of all relevant parameters and variables see Tab. I.

**TABLE I.** Overview of our parameters and observables.

|  |  |
| --- | --- |
| $L$ | template length |
| $k_0$ | basal extension rate |
| $t_0$ | basal extension timescale ($1/k_0$) |
| $\tau$ | time window for copying or cycle duration |
| $l$ | size of error cluster at the terminus |
| $k_l$ | extension rate following an error cluster of size $l$ |
| $\sigma_l$ | stalling factor after error cluster of size $l$ |
| $\epsilon_l$ | error probability after error cluster of size $l$ |
| $d$ | distance to next upstream error cluster |
| $l'$ | size of next upstream error cluster |
| $p(d, l')$ | probability of unbound terminus |
| $f_e(\tau)$ | error fraction |
| $Y(\tau)$ | yield of completed strands |
| $t_{\text{perf}}$ | average copying time of an error-free product |
| $k_{\text{on}}$ | primer-template association rate, set to $10^7 \text{ s}^{-1}$ |
| $c_{\text{prim}}$ | primer concentration |
| $\tau^*$ | optimal cycle duration |
| $E_{\text{waste}}$ | wasted energy per completed copy |
| $\Delta G_{\text{lg}}$ | activation energy per nucleotide |
| $t_{\text{cop}}(\tau)$ | average time for the first error-free copy to appear |
| $t_{\text{cop}}^*$ | minimum of $t_{\text{cop}}(\tau)$ with respect to $\tau$ |

### Kinetic separation of different error classes

Using the model of Fig. 2B, we simulate non-enzymatic template-directed polymerization with template length *L* = 20, tracking the number of full-length copies and their copying errors over time. We extract the time to completion for each full-length copy, to obtain the statistical distributions of completion times with the corresponding mean and median values for different error classes with a specified number of errors (Fig. 3A). Here and below, all shown results are for DNA parameters whereas the corresponding plots for RNA parameters are shown in the *Figure supplement*. As expected, copies containing no or few errors are completed much faster than highly erroneous copies. Error-free full-length copies display a near-Gaussian distribution of completion times (Fig. S1 in *Figure supplement*) peaked at the mean completion time for perfect copies, *t*_perf_ = *Lt*_0_ (inset of Fig. 3A).

**FIG. 3.**
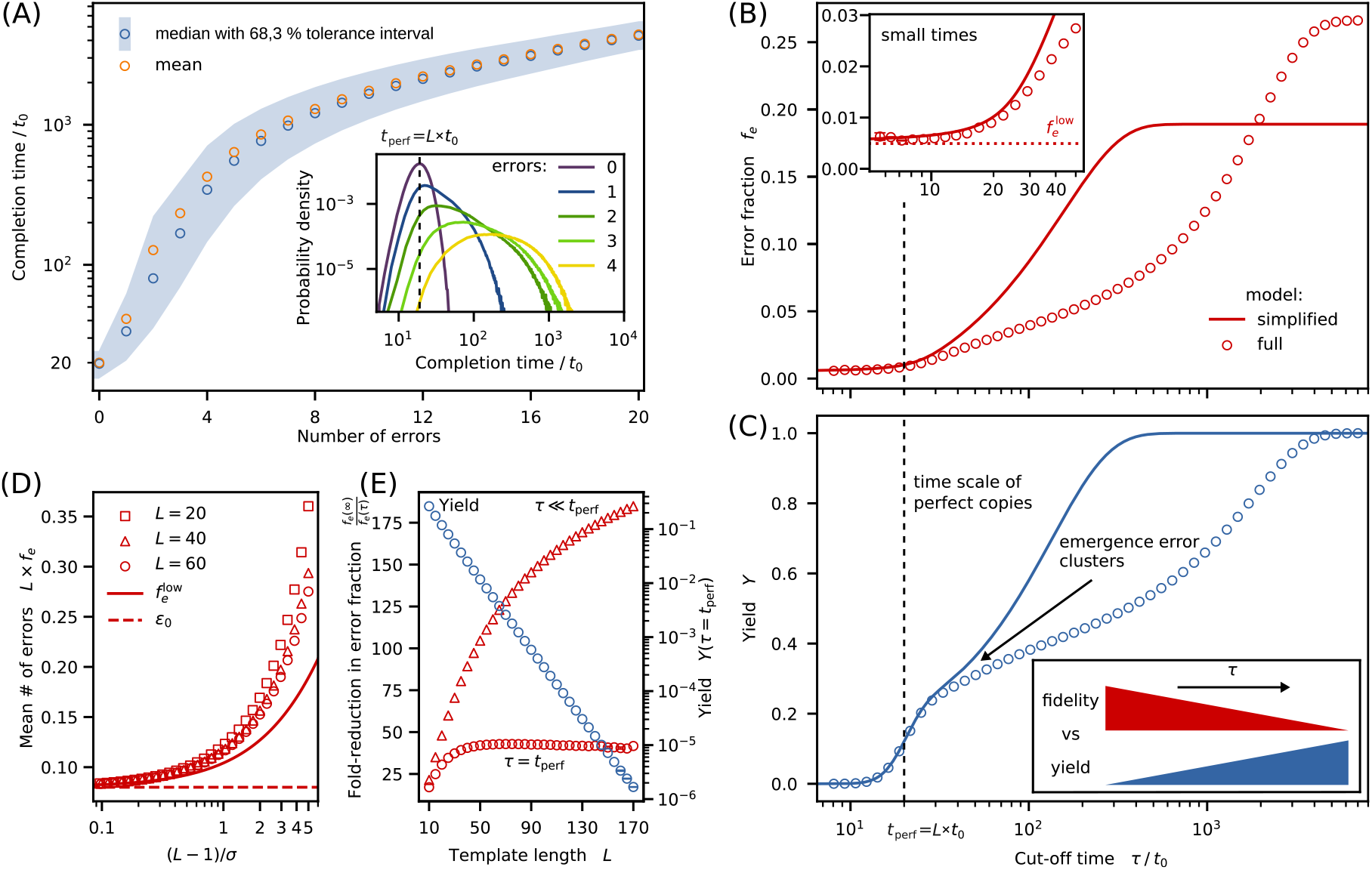
(A) Kinetic separation of different error classes. Typical copying times increase rapidly with the number of errors, as shown here for a template of length *L* = 20. Main panel: Mean and median completion times with centered 68.3 % confidence interval. Inset: Distributions of the completion time for zero to four errors (normalization such that the sum of areas below the curves for all possible error numbers is one). The zero-error distribution peaks at *t*_perf_ = *Lt*_0_, but overlaps with the distributions for one or more errors. (B) Fraction of errors in full-length copies, *f_e_*(*τ*), and (C) yield *Y*(*τ*) as a function of the copying time window *τ*. The increase in copying fidelity for smaller *τ* is at the expense of the yield. Lines show the analytical solution of the simplified model, whereas data points are obtained from stochastic simulations of the full model (error bars shown only for statistical errors > 1% of the mean). The simplified model approximates the full model well when *τ* is not much larger than the mean completion time of perfect copies. (D) In the short time limit (*τ* → 0), the absolute number of errors within completed copies depends only on (*L* – 1)/*σ*. For strong stalling (*σ* > *L*), the mean error number *Lf_e_* (symbols) is well approximated by the lower bound 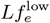 (solid line). For *σ* ≫ *L*, 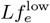 reduces to the error probability *ϵ*_0_. For a fixed value of (*L* – 1)/*σ* > 1, longer strands contain fewer errors than short ones. (E) The error fraction *f_e_* can be decreased significantly by reducing *τ* compared to the error fraction obtained for *τ* → ∞. The reduction of the error fraction achieved by choosing *τ* = *t*_perf_ goes hand in hand with a reduction of the yield *Y*.

Almost all copies, even the worst ones, reach full length within a completion time of *Lσ*_3_*t*_0_. Copies with few errors show asymmetric distributions of completion times (Fig. S1 in *Figure supplement*), with tails at small times that overlap with the error-free distribution (inset of Fig. 3A), which has implications for the limits of kinetic error discrimination (see below). We first turn to a trade-off that is inherent to kinetic error discrimination: Shortening the time window *τ* increases the fidelity of the obtained full-length copies, but decreases the yield.

### Fidelity-yield trade-off

We measure the fidelity of the copying process via the error fraction *f_e_*(*τ*), defined as the average fraction of wrongly incorporated nucleotides in full-length products when template-directed polymerization is stopped after time *τ*. The yield *Y*(*τ*) of the copying process is the fraction of templates for which copying has completed. Both, the error fraction and the yield increase with time and saturate when *τ* > *Lσ*_3_*t*_0_ (circle symbols in Figs. 3B and 3C). The yield approaches 100%, since our model does not contain any side reactions such as template cleavage by hydrolysis. However, the error fraction concomitantly reaches values larger than 0.25. This is clearly too high to conserve, e.g., the function of a ribozyme, even if the ribozyme is relatively robust against mutations [47]. In the other extreme of very short times *τ*, the error fraction is dramatically reduced to less than 1%, but the yield becomes essentially zero.

The vertical dashed lines in Figs. 3B and 3C mark the mean completion time *t*_perf_ of perfect copies. In this regime, the yield grows strongest while the error fraction still remains low. Hence, for a single copying cycle, values of *τ* ≈ *t*_perf_ represent the best compromise. However, another interesting feature of the fidelity curve in Fig. 3B is that *f_e_*(*τ*) apparently approaches a nonzero lower limit as *τ* → 0, which also appears consistent with the overlapping completion time distributions (inset of Fig. 3A). What factors determine this limit on how much kinetic error discrimination can improve the copying accuracy?

### Kinetic error discrimination is limited

To understand the error discrimination for short times, we turn to an analytically solvable simplified model, which accurately describes the behavior in this regime. The underlying approximations rely on the observation that copies containing a cluster of multiple consecutive errors have a negligible probability to be completed at short times *τ*. Hence, we ignore the dependence of the model parameters on the cluster size *l* by using a single stalling factor *σ* and a constant error probability *ϵ_l_* for all *l* > 0. Furthermore, we neglect the effect of preceding error clusters, i.e., we set *p*(*d*, *l*′) = 0, such that the copying rate is restored to *k*_0_ immediately after one correct incorporation. We derive the analytical expressions for the resulting error fraction and yield in *Appendix A*.

The analytical solutions for *σ* = *σ*_1_ (solid lines in Fig. 3B and 3C) confirm that the simplified model is equivalent to our full model for small times *τ*. They deviate when the first error clusters emerge: In the simplified model, (i) the yield grows more rapidly and saturates earlier, since error clusters do not increase stalling, and (ii) the error fraction is smaller, since the error probability does not grow after the first mismatch. For *τ* → ∞, the simplified model predicts an error fraction *ϵ*_0_/(1 + *ϵ*_0_ – *ϵ*_1_) in the limit of long templates (but with *L* fixed). This limit is independent of the stalling factor *σ*, since all strands reach full length.

Importantly, the simplified model lets us determine the lower limit 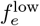 of the error fraction by including only copying processes with one or no error (see *Appendix A*),

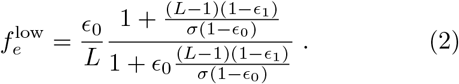

In the strong stalling regime (*σ* ≫ *L*) this reduces to 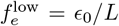, reflecting the absence of a kinetic penalty for errors in the last copying step. Fig. 3D compares the scaling of the lower bound (solid line) with (*L* – 1)/*σ* to the corresponding full analytical expression (symbols) in the limit *τ* → 0 for different values of *L*. For *L* < *σ*, the curves collapse onto one line and are well approximated by the lower bound, while they start to separate beyond this regime. Interestingly, long strands contain less errors than short ones for a fixed value of (*L* – 1)/*σ*.

Returning to the full model, we ask how much the error fraction can be lowered by reducing *τ*. We consider the fold-reduction in the error fraction, *f_e_*(∞)/*f_e_*(*τ*), evaluated both for *τ* ≪ *t*_perf_ and *τ* = *t*_perf_ at different template lengths (Fig. 3E). The first case merely illustrates the maximal possible fold-reduction at the cost of a vanishingly small yield. In contrast, the second case illustrates what is realistically attainable with a yield that is sizeable at small lengths, but decreases exponentially with *L* (Fig. 3E).

Why does the accuracy increase with length when *τ* ≪ *t*_perf_? The finite error fraction is mostly due to isolated errors, since error clusters would strongly stall the copying process and hence prevent the copy from reaching full length. However, isolated errors are rare since an initial mismatch is likely to trigger an error cascade. The longer the strand, the higher the probability for an error-cluster at some point. Long strands with isolated mismatches are thus more unlikely than short strands.

### Quantitative model for the kinetic error filtering scenario

We now turn to the full scenario for kinetic error filtering (Fig. 2A) and follow one template over a long observation time in a periodically changing environment. It is clear from the above analysis that short cycle times *τ* will lead to high accuracy, while the yield *Y*(*τ*) per cycle will be poor. However, the overall system output is determined by the yield rate, i.e., the yield per unit time, *Y*(*τ*)/*τ*. We assume that the duration of the temperature peak in the full scenario is much shorter than *τ*, and that the peak temperature is high enough to separate templates from both partial and completed copies. In addition to the template-directed polymerization process, we have to account for template-primer binding (Fig. 2A). Experiments [48–50] suggest an association rate *k*_on_ of about 10^7^ s^-1^ M^-1^. With an extension time of *t*_0_ = 1h, we have *k*_on_ = 3.6 × 10^10^/(*t*_0_ M). In our stochastic simulations, the time for each association event is drawn from an exponential distribution with mean 1/*k*_on_*c*_prim_, where *c*_prim_ is the primer concentration.

Since kinetic proofreading consumes free energy to increase fidelity, we also seek to analyze the unproductive free energy consumption of kinetic error filtering. Every time a covalent bond is formed a leaving group is consumed. However, the assembly of copies that remain incomplete at the end of a cycle is unproductive. Thus, the wasted free energy per completed copy is

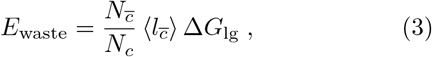

where *N_c_* is the number of completed copies produced during the observation time *t*, 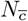 the number of incomplete copies, 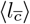 the average length of incomplete copies, and △*G*_lg_ the activation free energy per leaving group.

### Cyclic environments mitigate the fidelity-yield trade-off

We revisit the trade-off between fidelity and yield in our full scenario for kinetic error filtering. We explore the behavior over all possible cycle durations *τ*, since natural non-equilibrium environments display a broad range of timescales over which their physico-chemical conditions vary (e.g., due to convective cycles, as discussed above). Per cycle, the yield (Fig. 4A) and the error fraction (Fig. 4B) for a template of length *L* = 20 display essentially the same *τ*-dependence as observed before in Figs. 3B and 3C, except for an additional dependence on the primer concentration, which affects the timescale of template-primer binding. However, the more relevant quantity now is the yield rate *Y*(*τ*)/*τ*. Remarkably, the yield rate displays a peak as a function of *τ* (Fig. 4C). The peak becomes more pronounced with increasing primer concentration. At our largest concentration, *c*_prim_ = 1 nM, where the association time of the primer-template complex is negligible, the yield rate peaks at a cycle time close to *t*_perf_ (Fig. 4C).

**FIG. 4.**
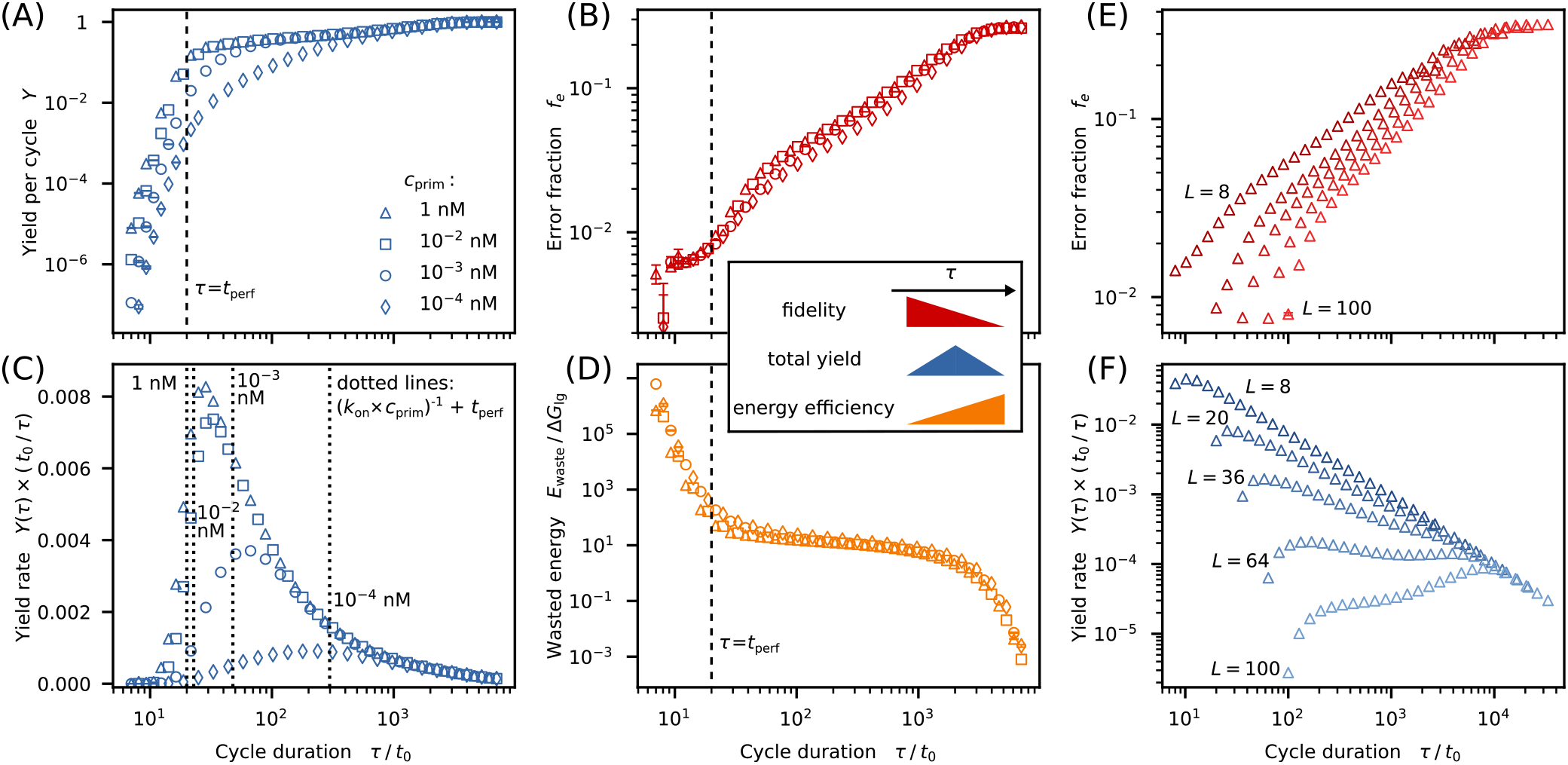
Fidelity-yield-energy trade-off in cyclic environments (kinetic error filtering). The (A) yield per cycle and (B) mean error fraction both decrease as *τ* is lowered (the vertical dashed line marks *t*_perf_). (C) However, the yield rate *Y*(*τ*)/*τ* displays a maximum at intermediate *τ* values (vertical dotted lines indicate the timescales (*k*_on_ *c*_prim_)^-1^ + *t*_perf_ for comparison), such that fidelity and yield can be increased simultaneously over a wide range of *τ* values. (D) Stronger error filtering is also coupled to an increased energy waste, which is proportional to the number of nucleotides contained in uncompleted copies. (E) Error fraction *f_e_*(*τ*) and (F) yield rate for different template lengths *L* (at fixed *c*_prim_ = 1 nM).

The peak in Fig. 4C implies that the fidelity-yield trade-off disappears over a certain range of cycle times: The fidelity and yield rate increase simultaneously as the cycle period *τ* is reduced from large times, until a value *τ*\* is reached where the yield rate is optimal. The *τ*-range of this simultaneous increase is largest for *c*_prim_ = 1 nM, whereas the effect becomes weaker for smaller concentrations.

The simultaneous increase of fidelity and yield rate comes at a free energy cost: The wasted free energy per completed copy, *E*_waste_, increases monotonically with decreasing *τ* (Fig. 4D). For *τ* ≈ *t*_perf_, hundred or more activated nucleotides are wasted per full-length copy, depending on the primer concentration. In constrast, for large *τ* almost no activated nucleotides are wasted.

How does the behavior in cyclic environments depend on the template length? Figs. 4E and 4F display the the error fraction *f_e_*(*τ*) and the yield rate *Y*(*τ*)/*τ* for different lengths *L* at the same primer concentration (*c*_prim_ = 1 nM). With increasing *L*, the yield maximum moves to longer cycle durations, becomes less pronounced, and eventually disappears. Concomitantly, a second maximum emerges at larger cycle periods, and hence larger error fractions. A significant increase in fidelity at high yield is only possible for lengths at which the left maximum still exists (see Fig. S2 in *Figure supplement*).

### Error-free copying

Is the fidelity-yield-boost in cyclic non-equilibrium environments sufficient to facilitate the emergence of molecular evolution? To not immediately lose a newly discovered functional sequence, a primitive copying process must at least form one accurate copy before the sequence is destroyed, e.g. by hydrolysis [15]. How long does it take to obtain the first error-free copy? To address this question, we compute the average copying time *t*_cop_, defined as the average time for the first perfect copy to appear. We then compare *t*_cop_ to typical lifetimes of template sequences.

Using the exact results for our simplified model (*Appendix A*), we derive an analytical expression for *t*_cop_ that is also valid for the full model (*Appendix B*). For *τ* ≫ (*k*_on_ *c*_prim_)^-1^, this expression simplifies to

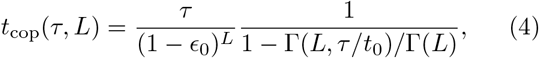

where Γ(*L*, *τ*) and Γ(*L*) are the incomplete and regular gamma function, respecitvely. Fig. 5A shows *t*_cop_ as a function of the cycle duration *τ* for different lengths at fixed *c*_prim_ = 1 nM. All curves exhibit a distinct minimum at an optimal cycle duration *τ*\*. The copying time 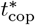 at the optimal cycle duration increases roughly exponentially with the template length (Fig. 5B and *Appendix B*), while *τ*\* grows roughly linearly with *L* (Fig. 5C). As long as *c*_prim_ ≥ 10^-2^ nM, the primer concentration does not significantly affect 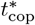. In this regime, and assuming *t*_0_ = 1 h, a first perfect copy of a 20-mer would typically arise within a few days, whereas about 20 weeks would be required for a 50-mer.

**FIG. 5.**
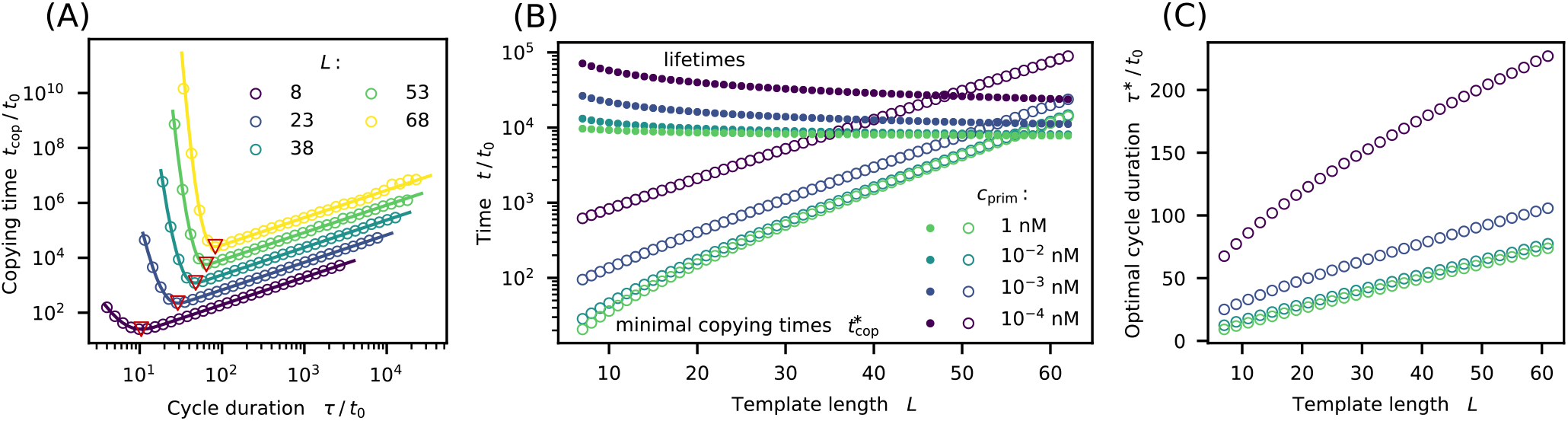
Kinetic error filtering at the minimal copying time optimum. (A) Copying time *t*_cop_ as a function of the cycle duration *τ* for different template lengths at *c*_prim_ = 1 nM. All curves display a distinct minimum (red triangles) at an optimal cycle duration *τ*\*. Symbols show data from stochastic simulations, while lines are obtained from Eq. B4. (B) The minimal copying time 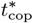 increases roughly exponentially with the template length, while the average template lifetimes decrease with *L*. These two timescales become equal for template lengths around 50 bases, depending on the primer concentration *c*_prim_. (C) Dependence of the optimal cycle duration *τ*\* on *L*.

How does the length-dependent optimal copying time 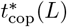 compare to the lifetime of the template? For temperatures below 37° C, DNA hydrolysis is hardly measurable, but the lifetimes decrease rapidly with temperature [51, 52]. In environments with temperature peaks that separate copies from templates, the high temperature phases limit the template lifetime [30]. To estimate lifetimes, we apply the same temperature profile and environmental conditions as in the experiment of Ref. [30] and use the predictive formula for degradation rates given in Ref. [51], see *Materials and methods* for details. The resulting lifetimes depend on *L* (Fig. 5B), since the number of hydrolysis sites increases with *L*, while the number of temperature cycles decreases with *L* due to the growing optimal cycle duration (Fig. 5C). The length where the copying time 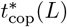 matches the estimated lifetime is around *L* ~ 50, with only a weak dependence on the primer concentration (Fig. 5B). Hence, kinetic error filtering can give rise to at least one error-free copy of DNA ~50-mers during their lifetime. With RNA parameters, this length threshold is at ~25-mers (see Fig. S6 in *Figure supplement*).

## DISCUSSION

The mechanistic basis of kinetic error discrimination in non-enzymatic copying of nucleic acid sequences is experimentally well documented: initial errors stall the copying process and increase the error probability for subsequent nucleotides [20, 21]. We showed that these molecular effects give rise to a strong kinetic discrimination against errors in full-length copies, if the time window for template-directed polymerization is sufficiently short (Fig. 3A). When this kinetic error discrimination mechanism is embedded in a length-selective cyclic environment, a kinetic error filtering scenario emerges (Fig. 2A) with several interesting features: (i) Kinetic error filtering does not require any sophisticated enzymes, and could thus act as a prebiotic precursor to kinetic proofreading in template-directed polymerization. (ii) Kinetic error discrimination displays an intrinsic fidelity-yield trade-off (Figs. 3B and 3C). This is in contrast to kinetic proofreading, which displays an intrinsic speed-accuracy trade-off [53]. However, the cyclic environment of the kinetic error filtering scenario creates a regime, in which the fidelity-yield trade-off is broken, such that reducing the cycle time *τ* simultaneously increases both, fidelity (Fig. 4B) and yield (Fig. 4C), at the cost of chemical energy (Fig. 4D). Energy efficiency is likely not a primary concern for early copying scenarios (but might become increasingly important as prebiotic living systems become more sophisticated and compete with each other). (iii) The cycle time *τ* can also be chosen to minimize the average time *t*_cop_ required to produce the first exact copy of a template (Fig. 5A), rather than to maximize the yield (Figs. 4C and 4F). Importantly, kinetic error filtering could sufficiently reduce *t*_cop_ to faithfully copy up to ‘50-mer templates within their lifetime (Fig. 5B).

Kinetic error filtering could spontaneously arise in hydrothermal systems: Convective cycles produce periodic variations of the temperature and other physico-chemical properties of the local environment, creating limited time windows for copying, combined with enhanced loss and degradation rates for short strands [30]. Since the geometries of natural hydrothermal systems vary over a broad range, one may expect a correspondingly broad range of convective cycle times, such that different systems will naturally sample the *τ* values required for different template lengths and optimization criteria. Length selection could result, e.g., from an interplay between convection and thermophoresis [31, 32], from accumulation on mineral surfaces [33], or retention within lipid vesicles [34].

We note that the kinetic error filtering scenario studied here is inherently different from a previously described effect of post-mismatch stalling on the error threshold in a mutation-selection model [20]. The latter model describes the competition between replicators in an environment with constant carrying capacity, a scenario that may arise at a later evolutionary stage than considered here. On a mathematical level, kinetic error filtering relies on fluctuations and rare events in an explictly timevarying environment. In contrast, the effect reported in Ref. [20] results from a shift in the balance between the opposing average forces of mutation and selection.

For the stage of prebiotic evolution considered here, a key issue is the maintenace of a functional nucleotide sequence that may arise by chance. The sequence might display a catalytic activity that exerts a positive feedback onto its own synthesis, or a negative feedback onto its own degradation. Initially, its catalytic activity is likely not strong enough to significantly boost its replication. However, such a fledgling ribozyme (or DNAzyme) could further evolve, if a weak replication process supports its maintenance against degradation [54]. The relevant threshold for maintenance is that the sequence gives rise to at least one error-free copy before it is degraded, e.g. by hydrolysis [15]. Our analysis suggests that kinetic error filtering can reach this threshold for DNA sequences of up to ~50 bases and RNA sequences of up to ~25 bases (Figs. 5B and Fig. S6 in *Figure supplement*). Short ribozymes and DNAzymes already display remarkable catalytic abilities, with DNAzymes not (clearly) inferior to ribozymes [55, 56]. In enzyme-free RNA copying, an important fraction of errors is due to G:U wobble paring [14, 57]. It is unclear whether the ribonucleotides available under prebiotic conditions correspond to the canonical ones found in extant living systems [58–60]. Alternative ribonucleotides not prone to wobble pairing could enable higher copying fidelities. For instance, replacing U with 2-thio-U significantly increases the fidelity in template-directed polymerization [61], such that RNA-based systems might reach similar error probabilities as DNA-based systems. Taken together, it appears that kinetic error filtering could push the enzyme-free copying of nucleic acid sequences via template-directed polymerization across an important threshold, facilitating the emergence and maintenance of sequences with catalytic functions.

## MATERIALS AND METHODS

### Recovery from error clusters

The extension rate *k_l_* and defined in Eq. (1) depends on the length *l* of the immediate error cluster at the extension site through the stalling factor *σ_l_*. The error probability *ϵ_l_* for the next incorporated nucleotide also depends on *l*. Moreover, effective extension rate and error probability also depend on the distance *d* to the preceding error cluster and its size *l*′ (see Fig. 2B) and we write

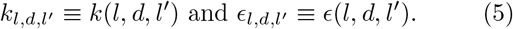

If a matching nucleotide is incorporated after a single mismatch, the extension rate recovers to an almost normal level according to experimental observations [21]. After two matches following the isolated error, no effect on the copying process was measured anymore. We therefore assume that a preceding error cluster of length *l*′ = 1 only affects the dynamics if *d* = 1. In this case, extension rate and error probability are given by *k*(*l*, 1, 1) = 0.67 × *k_l_* and *ϵ*(*l*, 1, 1) = 1.25 × for *ϵ_l_* ≤ 2.

In contrast, clusters of several mismatches influence the copying dynamics on a larger scale. The number of correctly incorporated nucleotides that are needed to restore the unperturbed extension dynamics increases with the size of the preceding error cluster. To include this effect, we use an extension of the binding model described in Ref. [21]. Matching nucleotides at the terminus following an error cluster are in a dangling configuration with probability *p*, which is a function of the distance d to and the size *l*′ of the preceding error cluster, i.e.,

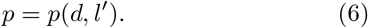

In Ref. [21], RNA folding software [62] is used to estimate p. It is shown, that the perturbation decays quickly with the number of correctly incorporated nucleotides. For *d* > 3, the probability to observe the terminus in a bound state is approximately one regardless of the value of *l*′ [21]. Within our model, the next copying steps after an error cluster are assumed to proceed regularly with *σ_l_* and *ϵ_l_* at probability 1 – *p*(*d*, *l*′). At probability *p*(*d*, *l*′) the copying process continues in a non-templated fashion with stalling factor and error probability given by *σ*_non_ and *ϵ*_non_. Probabilities for bound and unbound configurations are then accounted for in a coarse-grained fashion: Effective extension rate *k*(*l*, *d*, *l*′) and error probability *ϵ*(*l*, *d*, *l*′) are obtained by taking the average over the bound and dangling configuration, i.e.,

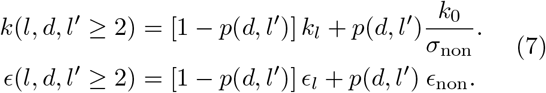

Numerical values for *p*(*d*, *l*′) used in the simulation are in line with Ref. [21] and are summarized in Tab. II.

**TABLE II.** A matching nucleotide at the primer’s 3’-end is in an unbound configuration with probability *p*(*d*, *l*′) where *d* and *l*′ denote the distance to and the size of the preceding error cluster.

| d | 1 | 2 | 3 | > 3 |
| --- | --- | --- | --- | --- |
| $p(d, l' \geq 3)$ | 0.9 | 0.5 | 0.1 | 0 |
| $p(d, l' = 2)$ | 0.5 | 0.1 | 0 | 0 |

### Analysis of fold-reduction in error fraction

The value for the error fraction in the limit *τ* ≪ *t*_perf_ in Fig. 3B and was obtained from the simplified analytical model. However, we assume that the analytical and the full stochastic model give similar results in the short time limit. Obtaining a data set suited for statistical analysis for *τ* < 0.5 *t*_perf_ from stochastic simulations was not possible since the yield was too poor for long templates.

### Estimate for lifetime of DNA and RNA strands

According to Ref. [51] the rate of hydrolysis for a single-stranded RNA oligomer of *L* bases at a temperature T can be predicted by the following empirical formula:

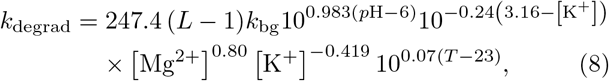

where *k*_bg_ = 1.3 × 10^-9^ min^-1^ is the background rate determined at *p*H 6, [K^+^] = 3.16 M and at 23° C. The temperature *T* in the last term on the right-hand side of Eq. (8) is expressed as a multiple of 1^°^ C.

To predict the lifetime of DNA and RNA strands, we assume environmental conditions similar to the experimental study performed with RNA polymers of length *L* = 60 in Ref. [30], i.e., [Mg^2+^] = 0.05M, [K^+^] = 0.05M and *p*H 8.3. Moreover, we assume the same profile for the temperature peaks separating copies from templates as in Ref. [30], i.e., *T* = 68° C for *τ*_hot_ = 5.56 × 10^-4^ h.

It is known that the stability of DNA strands against hydrolysis is much higher compared to RNA strands [51]. Therefore, we use the results based on Eq. (8) as a lower bound for the lifetime of DNA oligomers.

For temperatures below 37^°^ C, hydrolysis of DNA strands is hardly measurable [51, 52]. Hence, in the DNA scenario, we assume that hydrolysis only occurs during the temperature peaks. The average degradation rate over one optimal temperature cycle is then given by 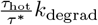.

In contrast to DNA strands, RNA strands are also prone to hydrolysis at lower temperatures. To estimate the lifetime of an RNA oligomer, one, therefore, has to compute an average rate of hydrolysis taking, into account the degradation rate during the cold phase 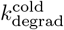 and the degradation rate during the temperature peaks *k*_degrad_. To obtain the correct average, 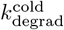 and *k*_degrad_ have to be weighted with the durations of the cold phase and the hot phase, respectively, i.e., 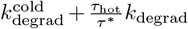. (Note that the duration of the cold phase is approximately equal to the duration of the cycle.)

Our estimates for the lifetimes are conservative, since hydrolysis within a double strand is rare compared to hydrolysis of a single strand. The double-strand configuration prevents the attacking 2’-OH from attacking the phosphodiester bond [52]. During the copying process, the template strand is partially protected by the extended primer. Therefore, the actual degradation rate might be smaller.

## ACKNOWLEDGMENTS

We thank Bernhard Altaner, Dieter Braun, Patrick Kudella, and Joachim Rosenberger for fruitful discussions. UG thanks the KITP in Santa Barbara for hospitality on an extended visit, during which part of this work was completed. This work was supported by the German Research Foundation (DFG) via the TRR 235 Emergence of Life (Project-ID 364653263) and the excellence cluster ORIGINS. This research was also supported in part by NSF Grant No. PHY-1748958, NIH Grant No. R25GM067110, and the Gordon and Betty Moore Foundation Grant No. 2919.02.

## DATA AVAILABILITY

The datasets that support the findings of this study are available from the corresponding author upon reasonable request.

## CODE AVAILABILITY

The computer codes used to generate the datasets, are available from the corresponding author upon reasonable request.

## Appendix A: Analytical solutions for simplified model

In the main text, we introduced a simplified copying model to discuss the error fraction’s bounds. Here we will develop the model in detail and derive analytic expressions for the yield and the error fraction.

The simplified model partially neglects the polymerization history and only takes into account the last incorporation. If the nucleotide at the 3’-end of the (partially) extended primer is a mismatch, the copying process is stalled by a constant factor regardless of the number of preceding errors, i.e. *σ*_*l*=1_ = *σ*_*l*>1_ = *σ*. In the same way, the probability for the next incorporated nucleotide to be a mismatch also remains constant beyond the first mutation, ie. *ϵ*_l=1_ = *ϵ*_*l*>1_. We further assume that the unperturbed dynamics are restored immediately after the first correct monomer is built-in and that the end of the partially extended primer is always bound to the template strands, i.e., *p*(*d*, *l*′) = 0 ∀ *d*, *l*′. Hence, error clusters do not affect the extension dynamics on distances longer than one.

The master equation [63] describing the dynamics can be stated as follows: Define *p*_*m*,*n*_(*t*) as the probability to have polymerized *m* steps with *n* mutations, but none of them in the last step. Moreover, introduce *q*_*m*,*n*_(*t*) as the probability to have made *m* steps with *n* mutations, one of them in the last step. With that, we can write down the following equation for *p*_*m*,*n*_(*t*) where we use the symbol *k*_0_ for the basic extension rate as in the main text.

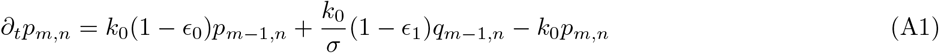

The first two terms on the right-hand side are gain terms and describe the extension of a strand of length *m* – 1 containing *n* mutations with a match following a match (first term) or following a mismatch (second term). The third one is loss term and accounts for the extension of a strand that has length *m* and contains *n* errors with either a match or a mismatch following a match. The equation for *q*_*m*, *n*_(*t*) reads as follows.

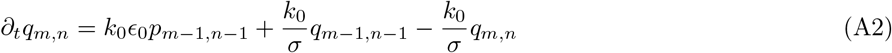

The two first terms on the right-hand side are the gain terms. The first one describes the extension of a strand of length *m* – 1 and *n* – 1 mutations with a mismatch following a match. The second term accounts for the stalled extension of a strand of the same length and same error number with a match or mismatch following a mismatch. The third term is a loss term for the stalled extension of a strand of length *m* with *n* mutations with a match or mismatch following a mismatch.

Laplace transformation of the above equations, such that 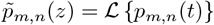, leads to

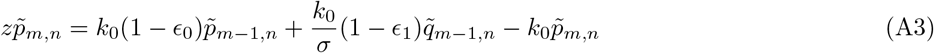

and to

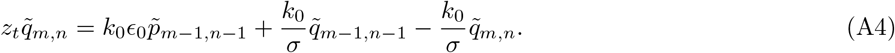

Some algebra suffices ot show that the solution is given by

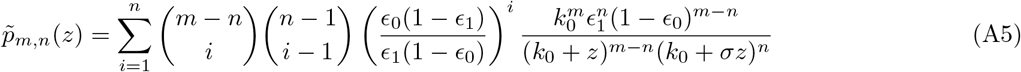

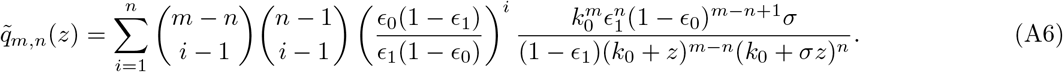

Backtransforming, we obtain

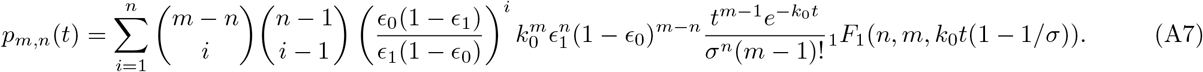

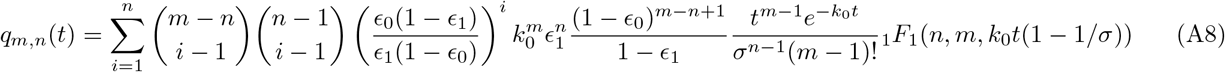

where _1_*F*_1_(*a*, *b*, *x*) is the confluent hypergeometric function [64] which can be written as

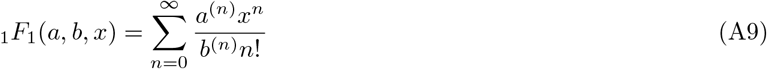

Here, *a*^(*n*)^ and *b*^(*n*)^ are rising factorials.

The relevant observable now is 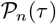, which denotes the probability to observe a complete polymerization product with *n* mutations when copying a template of length *L* after time *τ*. 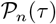 is given by

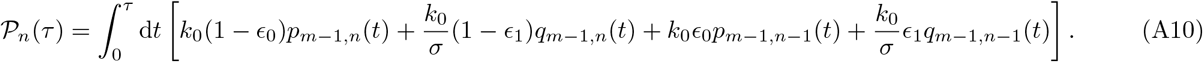

Introducing Φ_*L*,*n*_(*τ*) as

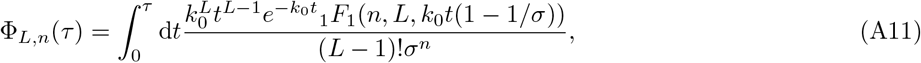

and using that

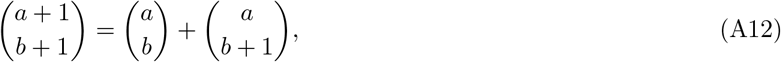

we can rewrite (A10) and obtain

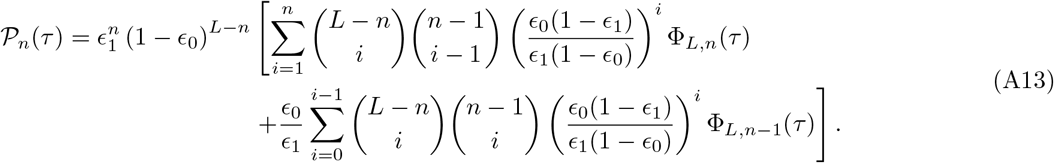

With Eq. (A13) error fraction and yield for copies of length *L* as a function of *τ* are then given by

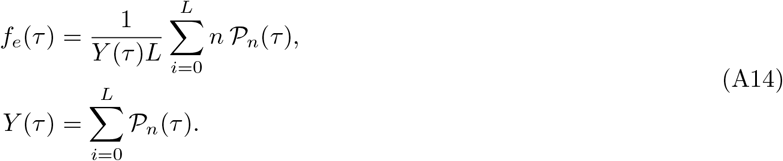

Eq. (A13) and (A14) have to be evaluated by numerical integration. In Fig. 6 *f_e_*(*τ*) and *Y*(*τ*) are plotted for *ϵ*_0_ = 0.08, *ϵ*_1_ = 0.69 and *σ* = 25 and compared to data from a corresponding stochastic simulation.

**FIG. 6.**
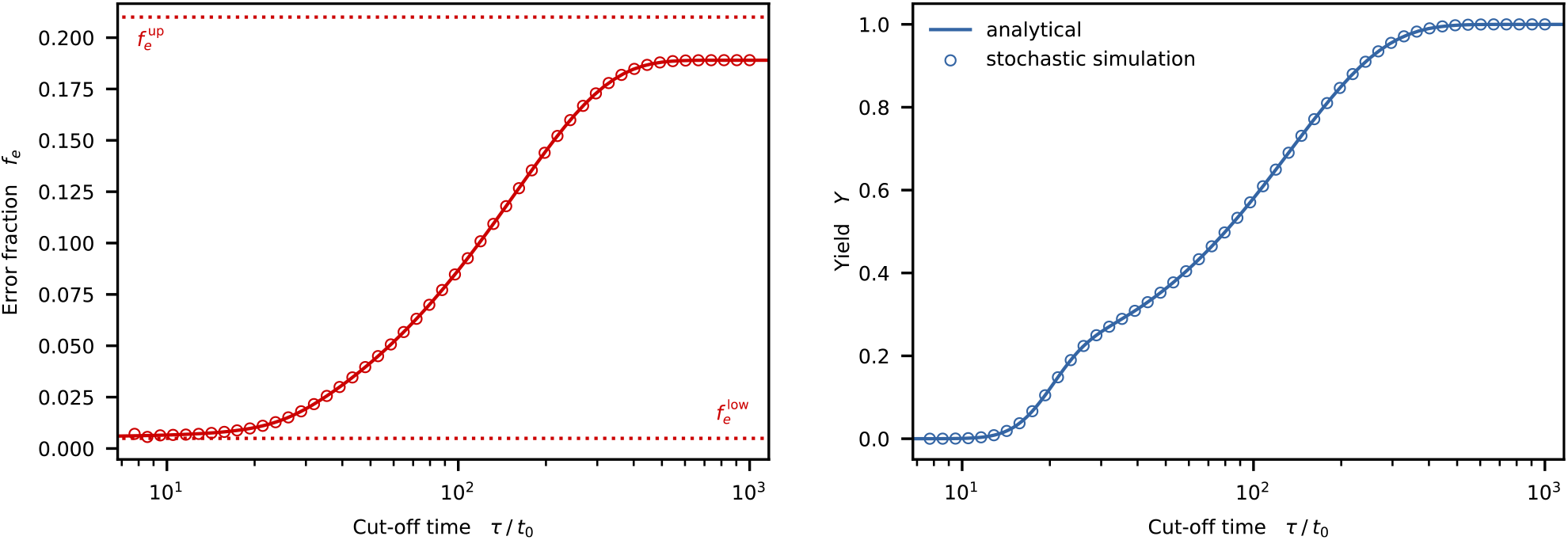
Error fraction *f_e_*(*τ*) and yield *Y*(*τ*) as functions of *τ* for *ϵ*_0_ = 0.08, *ϵ*_1_ = 0.69 and *σ* = 25 (see Eq. (A14)). Dotted lines indicate lower and upper bound for the error fraction according to Eq. (A17) and Eq. (A23). Circles: stochastic simulation (one data set).

In the limit *τ* → 0, an approximativ but compact analytical expression for the error fraction *f_e_*(*τ*) can be derived. In this limit, Eq. (A11) can be approximated as

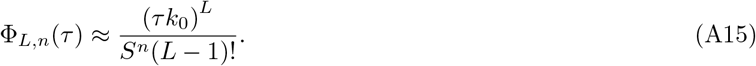

To arrive at Eq. (A15) the Taylor expansion of the integrand of Eq. (A11) is truncated at lowest order. From Eq. (A15) we also see that the yield goes to zero in this limit. For *τ* → 0 mostly copies containing no or only one mutation contribute to the yield. A lower bound for the error fraction in the short time limit can therefore be obtained as

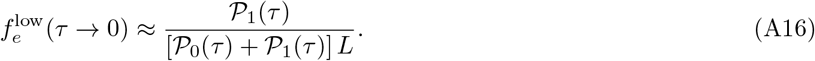

Plugging in Eq. (A15) into Eq. (A13) then transforms Eq. (A16) to

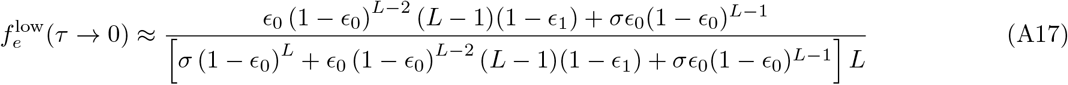

or to the more compact form used in the main text

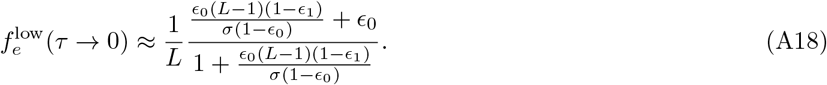

For the opposite limit *τ* → ∞ and for large *L* we can estimate the error fraction, which turns out to be an upper bound. In this limit, Eq. (A11) reduces to

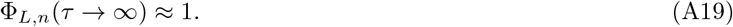

For long copies, it is justified to neglect the second term on the right-hand side of Eq. (A13) which corresponds to a copying trajectory with a misincorporation in the final step. Using that

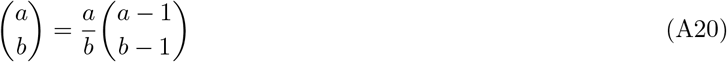

Eq. (A13) then becomes

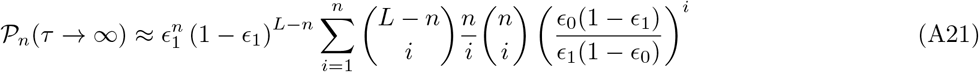

For *L* ≫ 1 this approximate expression for 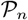 is sharply peaked. Neglecting the factor 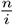 therefore only changes the overall scaling, which we will account for by the appropriate normalization later on. Replacing the binomials by their Gaussian approximations in Eq. (A21) then gives:

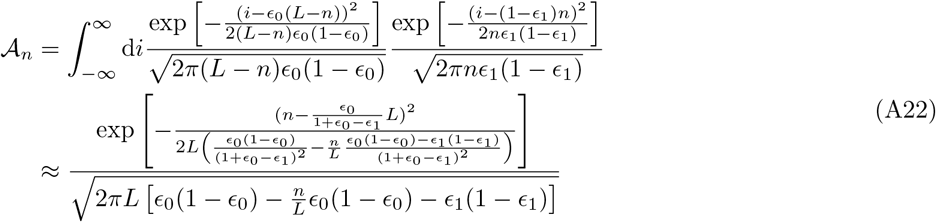

If we now replace 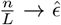 with some irrelevant but constant value in the denominators of the variance terms, we can give an expression for the error fraction:

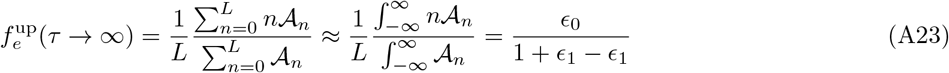

If *L* ≫ 1 templates almost have no chance to complete without any error. An initial misincorporation, in turn, is likely to trigger an error cascade. However, if templates are relatively short, there is a fair chance of not having an initial mismatch at all and therefore not running into an error cascade. Hence, we expect Eq. (A23) to be an upper bound for templates of finite length *L*.

## Appendix B: Analysis of the copying time

In the main text, *t*_cop_(*τ*, *L*) was introduced as the average waiting time for the first error-free copy to occur as a function of the cycle duration *τ* and the template length *L*. The differences in the dynamics between the simplified model (see *Appendix A*) and the full model only become apparent after the first mismatch got incorporated. For an error-free copying process leading to a full-length product, the dynamics are identical in both models. Hence, we can build on the analytic results obtained in *Appendix A* to derive a formula for *t*_cop_(*τ*, *L*).

According to Eq.(A13) the probability 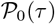 to observe a complete polymerization product without mutations after time *τ* is given by

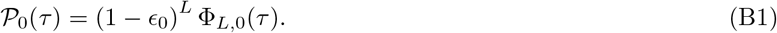

with

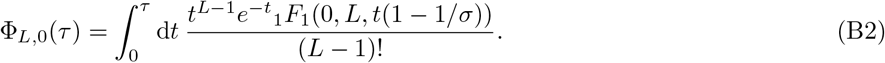

Note that in Eq. (B2) all timescale are expressed in units of the basal extension timescale *t*_0_ = 1h as in the main text. We will use this convention throughout this section. Using that the confluent hypergeometric function is identical to one if the first argument is zero [64], i.e., *F*_1_(0, *L*, *t*(1 – 1/*σ*)) ≡ 1, we obtain

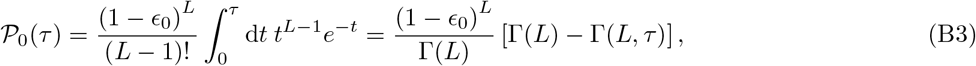

where Γ(*L*) and Γ(*L*, *τ*) are the gamma and upper incomplete gamma function.

In the cycling scenario *τ* corresponds to the cycle duration. If we assume, that the average association time 〈*t_a_*〉 = (*k*_on_ × *c*_prim_)^-1^ is short in comparison to the cycle duration *τ*, 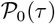 represents the probability to obtain an error-free product within one cycle. Then, the average number of cycles, one has to wait until the first error-free full copy is observed, is 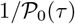 and the copying time *t*_cop_ (*τ*, *L*) is given as

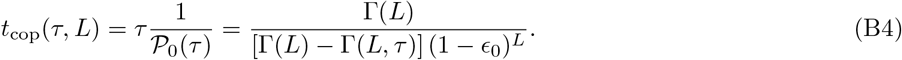

The situation is more complex if the primer concentration is small such that the association time *t*_a_ is not negligible anymore. In this case, the effective time window for the copying process within one cycle is given by *τ* – *t*_a_. *t*_a_ in turn is distributed exponentially. The probability of observing an error-free product at the end of the cycle, therefore, takes the form

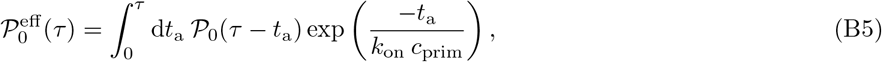

where the exponential on the right-hand side corresponds to the association time distribution. Carrying out the integral, taking the inverse, and multiplying with the cycle duration *τ* then leads to

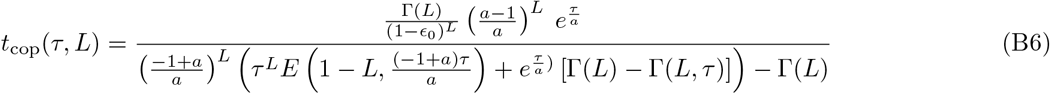

where we have used the abbreviation *a* = *k*_on_ *c*_prim_ and where *E*(*x*, *y*) is the exponential integral function which is defined as

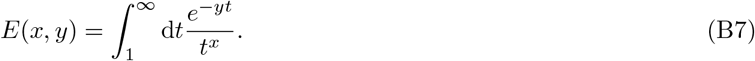

In Fig. 7 *t*_cop_(*τ*, *L*) according to Eq. (B6) is plotted for *ϵ*_0_ = 0.08 and *c*_prim_ = 10^-4^ nM for different values of *L* (straight lines) together with data points obtained from a corresponding stochastic simulation (circles). All curves show a distinct minimum. In general, the exact position of the minimum has to be determined numerically.

**FIG. 7.**
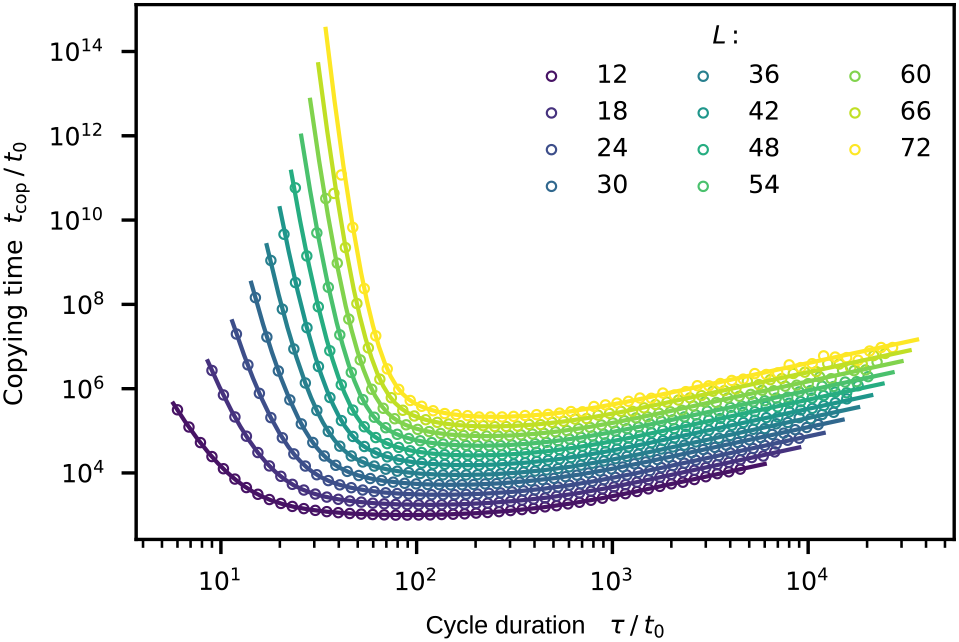
Copying time *t*_cop_ as a function of the cycle duration *τ* for *c*_prim_ = 10^-4^ nM. *t*_cop_ is the average time that passes until the first error-free copy of the template strand is produced. All curves show a distinct minimum. As the length *L* increases, the minimum moves to the right. Straight lines are obtained from Eq. (B6), circles are obtained from simulation (one data set).

However, for large primer concentrations, such that the association time becomes negligible, we can derive approximative formulas for the position *τ*\* and the value 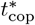 of the minimum of the copying time according to Eq. B4. From 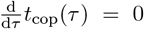, we obtain

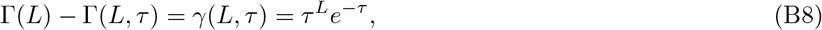

where *γ*(*L*, *τ*) is the lower incomplete gamma function defined as

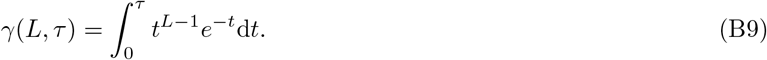

To be able to solve for *τ*, we first need an approximate expression for the value of *γ*(*L*, *τ*). The integrand *I*(*L*, *t*) in Eq. B9 can be written as:

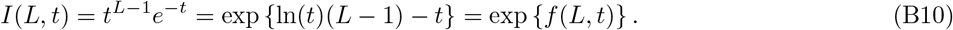

The value of *t* maximizing *f*(*L*, *t*) also maximizes I(L, t). From 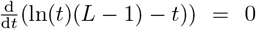 this value is determined to be *t* = *L* – 1. If we assume an upper integration boundary *τ* > *L* on the right-hand side of Eq. B9 we can perform a saddle point approximation to get an estimate for *γ*(*L*, *τ*). The assumption that *τ* > *L* is based on the observation that the yield per cycle drops quickly for *τ* < *L* (see Fig. 5C and Fig. 6A). According to the saddle point approximation, *γ*(*L*, *τ*) can be estimated as

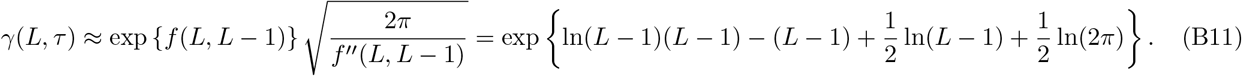

For *L* sufficiently large, we can neglect the last two terms in the exponential on the right-hand side and further replace (*L* – 1) by *L*. Plugging this simplified result into Eq. (B8) we obtain an equation determining the minimum position:

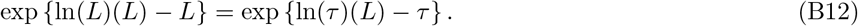

The position of the minimum is roughly given by *τ*\* ≈ *L*. In fact, *τ*\* > *L* since we neglected terms increasing *τ*\* (consistent with the saddle point approximation). Plugging this result into Eq. (B4) we obtain an approximative formula for the minimal copying time 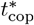:

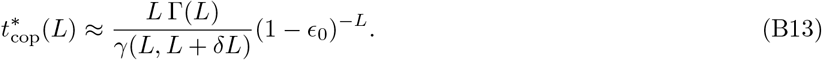

We wrote *L* + *δL* instead of *L* for the second argument of the lower incomplete gamma function in the denominator to emphasize that the upper integration boundary is actually slightly larger than *L*. Using Stirling’s approximation, i.e., 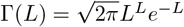 and the saddle point estimate for *γ*(*L*, *L* + *δL*) for *L* sufficiently large, we end up with simple expression for the minimal copying time as a function of L, i.e.,

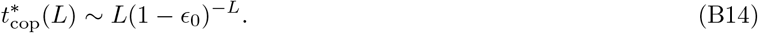

Hence, the scaling of 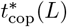 is dominated by the exponential factor exp {– ln(1 – *ϵ*_0_)}. In a plot with a logarithmic *y*-axis (decadic logarithm) this corresponds to a slope of ~ 0.036 for *ϵ*_0_ = 0.08. This result fits the slope observed in Fig. 5B.

