## Supplemental Figures for "A kinetic error filtering mechanism for enzyme-free copying of nucleic acid sequences"

(Dated: August 5, 2021)

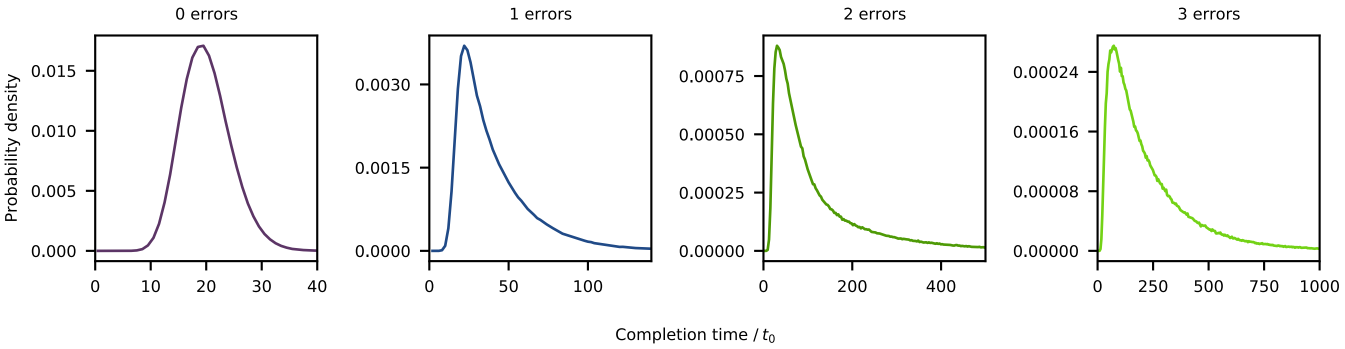


FIG. S1. Distributions of the completion time for zero to three errors. Typical completion times increase rapidly with the number of errors. The zero-error distribution approximately has the shape of a Gaussian distribution with a distinct peak at $t_{\mathrm{perf}} = Lt_0 = 20$.

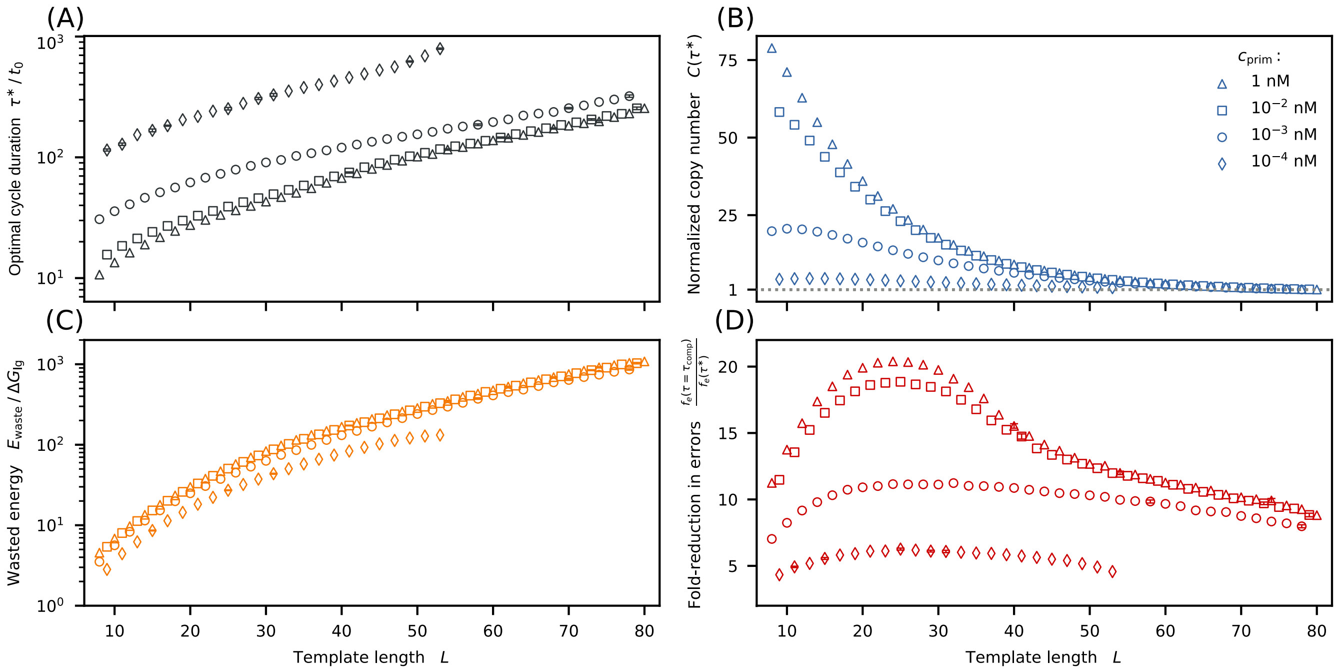


FIG. S2. Kinetic error filtering at the yield rate optimum. For a more systematic analysis of the effect of the template length, we introduce two new quantities: the cycle duration $\tau_{\text{comp}}$, for which 99% of all copies reach completion ($Y(\tau_{\text{comp}}) = 0.99$), and the ratio $C(\tau)$ of the yield rates evaluated at $\tau$ and $\tau_{\text{comp}}$, i.e., $C(\tau) = [Y(\tau) \times \tau_{\text{comp}}] \, / \, [0.99 \times \tau]$ . We refer to $C(\tau)$ as the normalized copy number, since it corresponds to the overall number of full copies obtained over a fixed time interval $\Delta t \gg \tau$ for a given cycle duration $\tau$ relative to the total number of full copies obtained in a scenario where the cycle duration is $\tau_{\text{comp}}$. The maxima of the normalized copy numbers for different template lengths $L$ are at the same positions as the maxima of the yield rates in Fig. 4F. With that, we can define "simultaneous optimization": A simultaneous optimization of fidelity and yield is possible for those values of $L$ for which the a left maximum of $C(\tau)$ exits and is larger than one. (A) Over the range of template lengths for which a simultaneous optimum exists, the optimal cycle duration $\tau^*$ increases with $L$. (B) The normalized copy number $C(\tau^*)$ decreases with $L$. As expected, smaller primer concentrations lead to smaller copy numbers. For low concentrations, as well as for larger $L$, the total yield is only slightly improved. (C) The wasted energy per completed copy evaluated at $\tau^*$ grows significantly as $L$ is increased. (D) The fold-change in the error fraction, $f_e(\tau_{\text{comp}})/f_e(\tau^*)$ is largest for template lengths between 20 and 30. However, the accuracy is significantly improved for all concentrations and lengths.

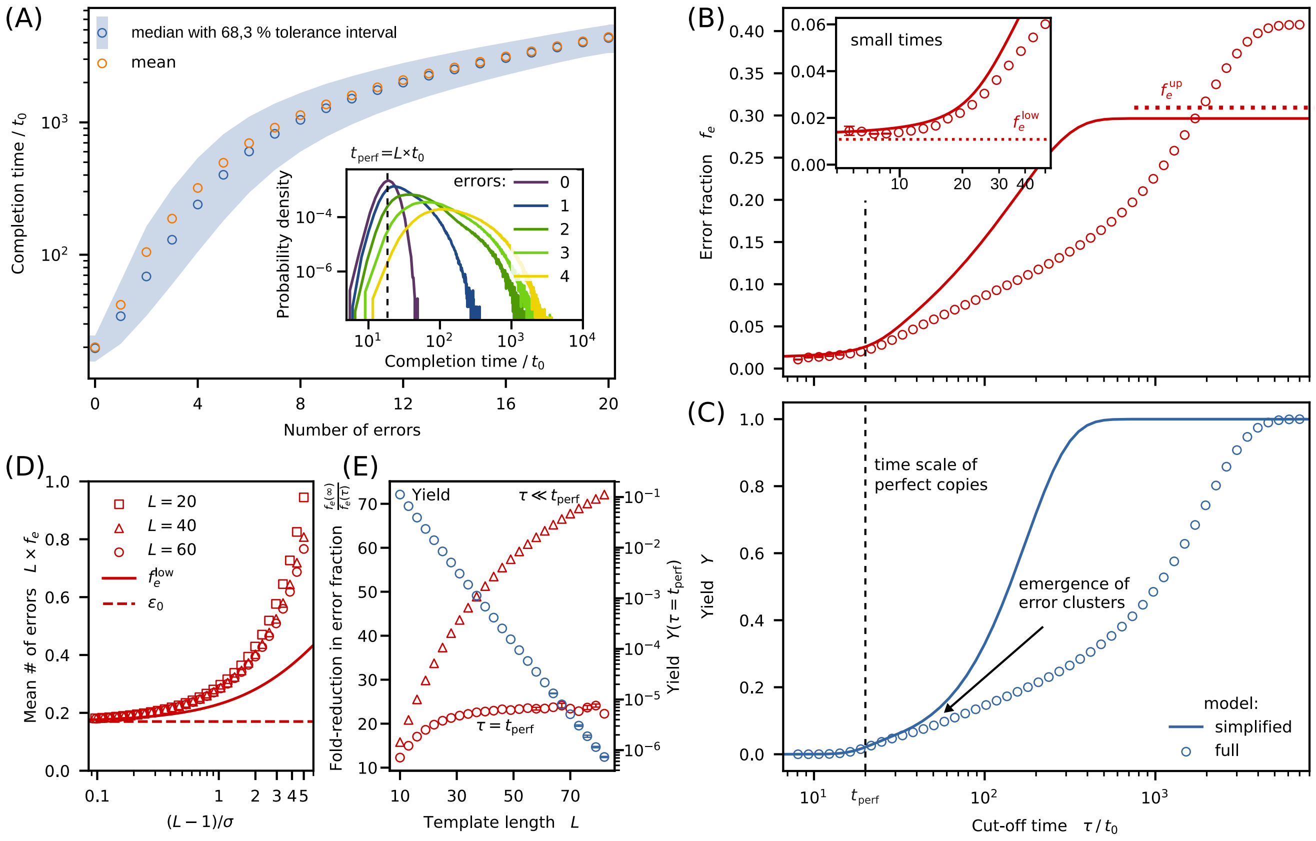


FIG. S3. Same as for Fig. 3 in the main text but with RNA parameters. (A) Comparing the probability densities for the RNA system to the DNA system shows that full-length products containing zero or only a few errors are more unlikely in the RNA system. (B) and (C) The error fraction for $\tau = \tau_{\mathrm{comp}}$ is more than a factor of 1.5 larger in the RNA system. Compared to the DNA system, the increase at $\tau = L\, t_0$ is less pronounced in the RNA system. Dotted red lines indicate the lower and upper bounds for the error fraction derived for the simplified model. (D) The absolute number of error for a given configuration in the RNA system is always larger than in the DNA system. (E) The error-reduction effect is by a factor of roughly tow smaller compared to the DNA system. The decrease of the yield is steeper in the RNA system (roughly a factor of two in the logarithmic plot).

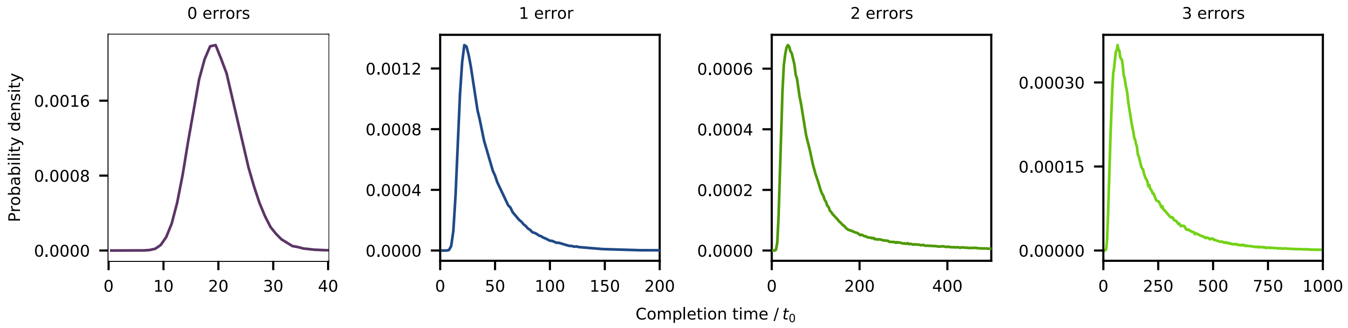


FIG. S4. Same as for Fig. S1 but with RNA parameters.

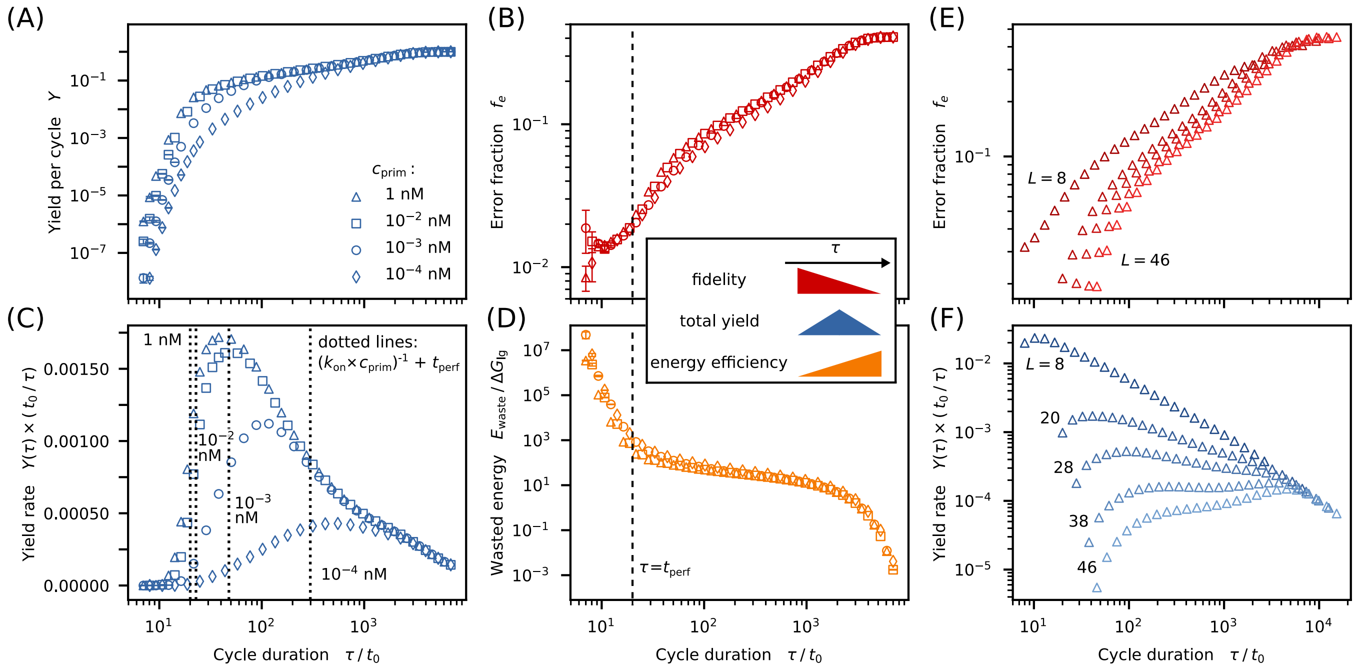


FIG. S5. Same as for Fig. 4 in the main text but with RNA parameters.

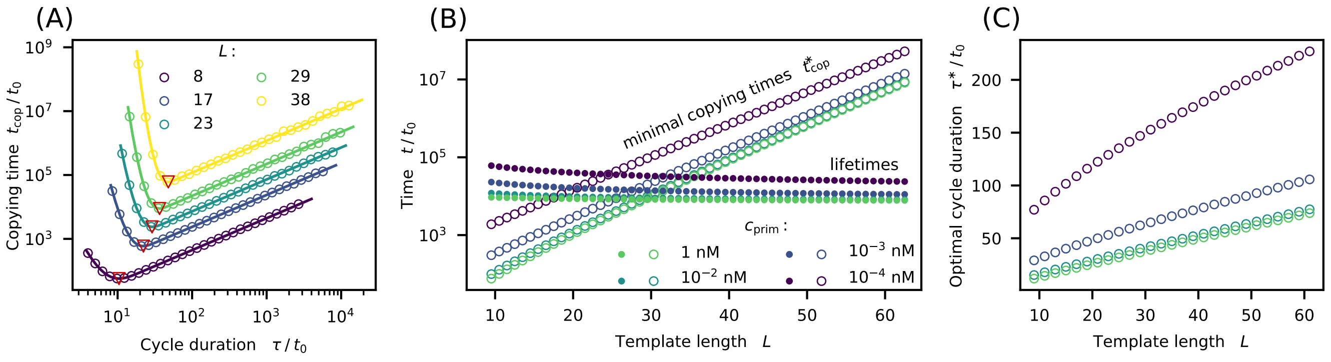


FIG. S6. Same as for Fig. 5 in the main text but with RNA parameters.

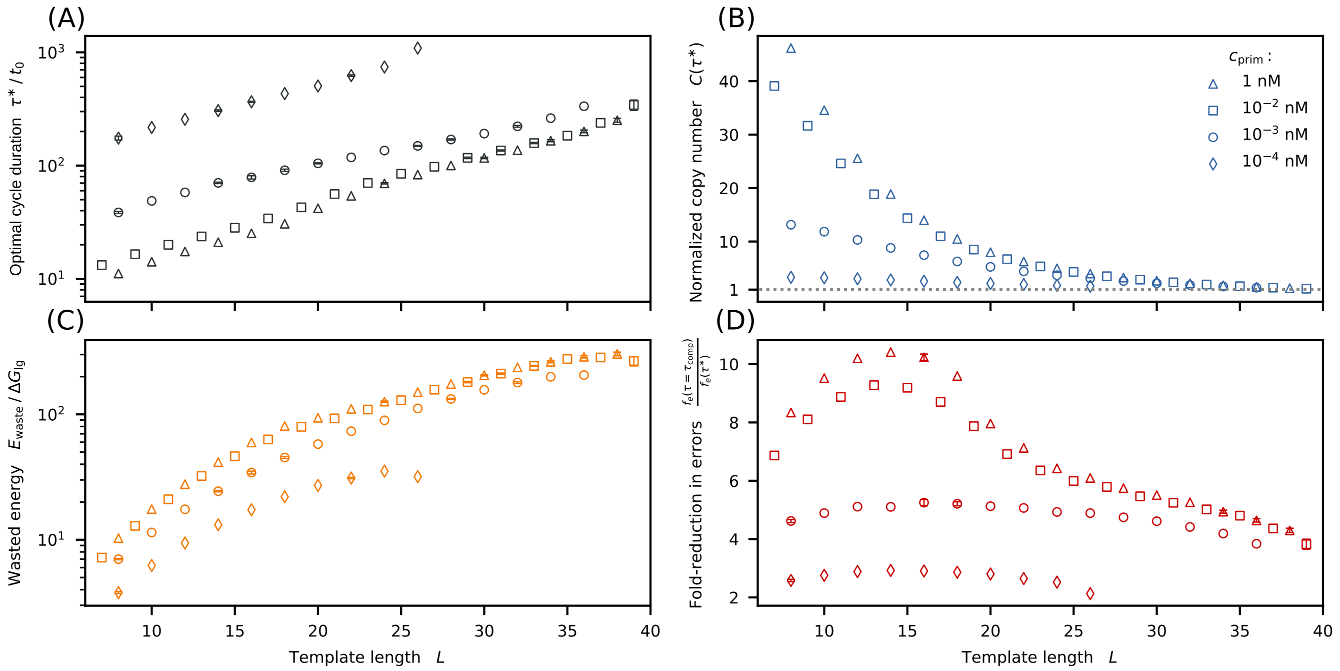


FIG. S7. Same as for Fig. S2 but with RNA parameters.
